# Learning modifies attention during bumblebee visual search

**DOI:** 10.1101/2023.10.05.561037

**Authors:** Théo Robert, Karolina Tarapata, Vivek Nityananda

**Affiliations:** Biosciences Institute, Newcastle University, Henry Wellcome Building, Framlington Place, Newcastle Upon Tyne, NE2 4HH, UK

**Keywords:** Reward, Saliency, Top-down attention, Insects, Bees, Cognitive Ecology

## Abstract

The role of visual search during bee foraging is relatively understudied compared to the choices made by bees. As bees learn about rewards, we predicted that visual search would be modified to prioritise rewarding flowers. To test this, we ran an experiment testing how bee search differs in the initial and later part of training as they learn about flowers with either higher-or lower-quality rewards. We then ran an experiment to see how this prior training with reward influences their search on a subsequent task with different flowers. We used the time spent inspecting flowers as a measure of attention and found that learning increased attention to rewards and away from distractors. Higher quality rewards led to decreased attention to non-flower regions, but lower quality rewards did not. Prior experience of lower rewards also led to more attention to higher rewards compared to distractors and non-flower regions. Our results suggest that flowers would elicit differences in bee search behaviour depending on the sugar content of their nectar. They also demonstrate the utility of studying visual search and have important implications for understanding the pollination ecology of flowers with different qualities of reward.

**Significance Statement:** Studies investigating how foraging bees learn about reward typically focus on the choices made by the bees. How bees deploy attention and visual search during foraging is less well studied. We analysed flight videos to characterise visual search as bees learn which flowers are rewarding. We found that learning increases the focus of bees on flower regions. We also found that the quality of the reward a flower offers influences how much bees search in non-flower areas. This means that a flower with lower reward attracts less focussed foraging compared to one with a higher reward. Since flowers do differ in floral reward, this has important implications for how focussed pollinators will be on different flowers. Our approach of looking at search behaviour and attention thus advances our understanding of the cognitive ecology of pollination.

## Introduction

Foraging bees must learn which flowers are rewarding and which ones are not. Given this ecological demand, they have evolved to be expert learners and are well-studied as models of visual cognition (Avarguès-Weber et al. 2011; Giurfa 2012). Bees learn to choose rewarding flowers and avoid differently coloured flowers without rewards (Lubbock 1881; Turner 1910; von Frisch 1914; Benard et al. 2006; Avarguès-Weber and Giurfa 2014). They are also capable of discriminating between higher and lower rewarding flowers (Baude et al. 2011; Riveros and Gronenberg 2012; Avarguès-Weber et al. 2018; Solvi et al. 2022). While a large body of research has demonstrated reward-based learning in bees, most of the work has looked at how learning affects the choices made by bees. Much less research has investigated the influence of rewards on visual search and attention in bees (Spaethe et al. 2006; Morawetz and Spaethe 2012; Nityananda and Pattrick 2013; Nityananda and Chittka 2021).

Visuospatial attention has been defined as a spotlight focussing on one region compared to others (Posner 1980) and is often measured by responses to targets in a region or the time spent looking at specific regions or objects (Schütz et al. 2011; Henderson and Hayes 2018). Visual search experiments look at how attention is deployed when searching for one target amongst others (Horowitz and Wolfe 2001). This approach has been used in several animals including jays, owls and fish (Dukas and Kamil 2000; Bond and Kamil 2002; Ben-Tov et al. 2015; Orlowski et al. 2015, 2018; Saban et al. 2017). Recent work has begun to look at attention and visual search in insects (Nityananda 2016), especially in bumblebees. Bees have been shown to flexibly switch between multiple rewarding targets (Nityananda and Pattrick 2013; Li et al. 2017). In experimental set-ups, floral rewards influence not just their choices but their visual attention, as measured by the time spent around particular flowers (Nityananda and Chittka 2021). Bees spend more time around higher rewarding flowers even when they are less salient than lower rewarding flowers. We still, however, know little about how bee visual search changes over time as the bees learn about rewards.

Bee attention during learning could be influenced by multiple factors, including the reward value and the saliency of the flowers. These factors have been shown to influence both bee choices and visual search. Colour contrast against a background, one measure of saliency, influences their visual search (Goulson 2000). Naïve bees also have an innate bias toward colours in the blue-green wavelength range and colours that have spectral purity (Lunau 1990; Lunau et al. 1996). We would therefore expect that the visual search of naïve bees would initially be influenced by saliency and innate biases. Subsequently, as bees learn about the reward value of flowers, we should expect them to pay more attention to rewarding flowers. We would also predict that there would be different effects on bumblebee visual search if the rewarding flowers had lower rewards or higher rewards. Given the effects of both reward value and stimulus saliency we would therefore expect learning to increased attention to higher reward but lower saliency flowers compared to lower reward high saliency flowers.

To test these ideas, we investigated the training bouts for bees trained as part of a previously published study (Nityananda and Chittka 2021) that focussed on the behaviour of bees in tests after the training. As part of that study, bees were trained on one of two flower types – either higher reward lower saliency flowers or lower reward higher saliency flowers. In both training regimes, the rewarding flowers were presented simultaneously with non-rewarding distractors. In the current study, we focus on the training period prior to the tests, that have not been previously analysed. We investigated the effect of learning on attention by comparing visual search in the initial period of the training with visual search in the final stage of the training. Prior expectations can change the perception of reward in social insects (Bitterman 1975; Gil et al. 2007; Wendt et al. 2019). In a second experiment, we therefore also investigated how prior experience of higher rewards and lower rewards influenced visual search when encountering new flowers that had different reward values.

We hypothesised that bees would increase their attention to rewarding flowers as they learnt about the rewards and that this effect would be greater for flowers with higher reward. We therefore predicted that in the first experiment, bees would attend more to rewarding flowers in the final phase of their training compared to the initial phase of their training. We further predicted that this change would be greater for the higher reward lower saliency flowers. Given the possibility of prior expectations influencing behaviour, we hypothesised that bees that had experienced higher reward should be less motivated by lower rewards. In the second experiment, we therefore predicted that bees would spend more attention away from rewarding flowers if they encountered lower rewarding flowers after having prior experience of higher rewards. We also predicted that bees would increase attention to rewarding flowers if the bees encountered higher rewards after experiencing lower rewards first.

## Materials and methods

### Bees

We obtained the bees from a commercial supplier (Syngenta Bioline, Weert, The Netherlands). We then tagged them with Opalith number tags (Christian Graze KG, Weinstadt-Endersbach, Germany) which allowed us to individually identify them. We transferred the bees under red light to one of two chambers of a wooden nest box (length × width × height: 28 × 16 × 11 cm). The floor of the other chamber was covered with cat litter to allow bees to discard refuse. This nest box was connected to an arena with a 24.5 cm long transparent Perspex tunnel. The arena consisted of a wooden box (length × width × height: 100 × 60 × 40 cm) covered with a UV-transparent Plexiglas lid and the arena floor was covered with green card. The arena was lit from above with two twin lamps (TMS 24 F with HF-B 236 TLD (4.3 kHz) ballasts; Philips, The Netherlands) fitted with Activa daylight full spectrum fluorescent tubes (Sylvania, New Haven, UK). Bees were allowed to forage for sucrose solution in the arena and provided ∼ 3g pollen directly in their colony every alternate evening.

### Spectral reflectance of flowers

We used an Avantes AvaSpec 2048 spectrophotometer (Anglia Instruments Limited, Soham, UK) along with a deuterium-halogen light source relative to a BaSO_4_ white standard to measure the reflectance spectra of the artificial flowers. We converted the spectra obtained into a bee-specific colour space (Chittka 1992) using the spectral sensitivity of bumblebee photoreceptors (Skorupski et al. 2007), the spectral distribution of the lights used and the spectral reflectance of the background. The colour hexagon space has three vertices representing the points of maximum excitation of the blue, green and ultraviolet (UV) photoreceptors of the bee (Figure 1A). The other three vertices correspond to the response to mixtures of approximately equal excitation of each combination of two photoreceptors. The Euclidean distance between the centre of the hexagon and each vertex is 1 and colour distance greater than 0.1 can be distinguished by bees without special training procedures. After plotting the reflectance values of our flowers in this space, we were able to measure the distance in perceptual space between them. These data are provided in the previous paper (Nityananda and Chittka 2021).

**Figure 1.**
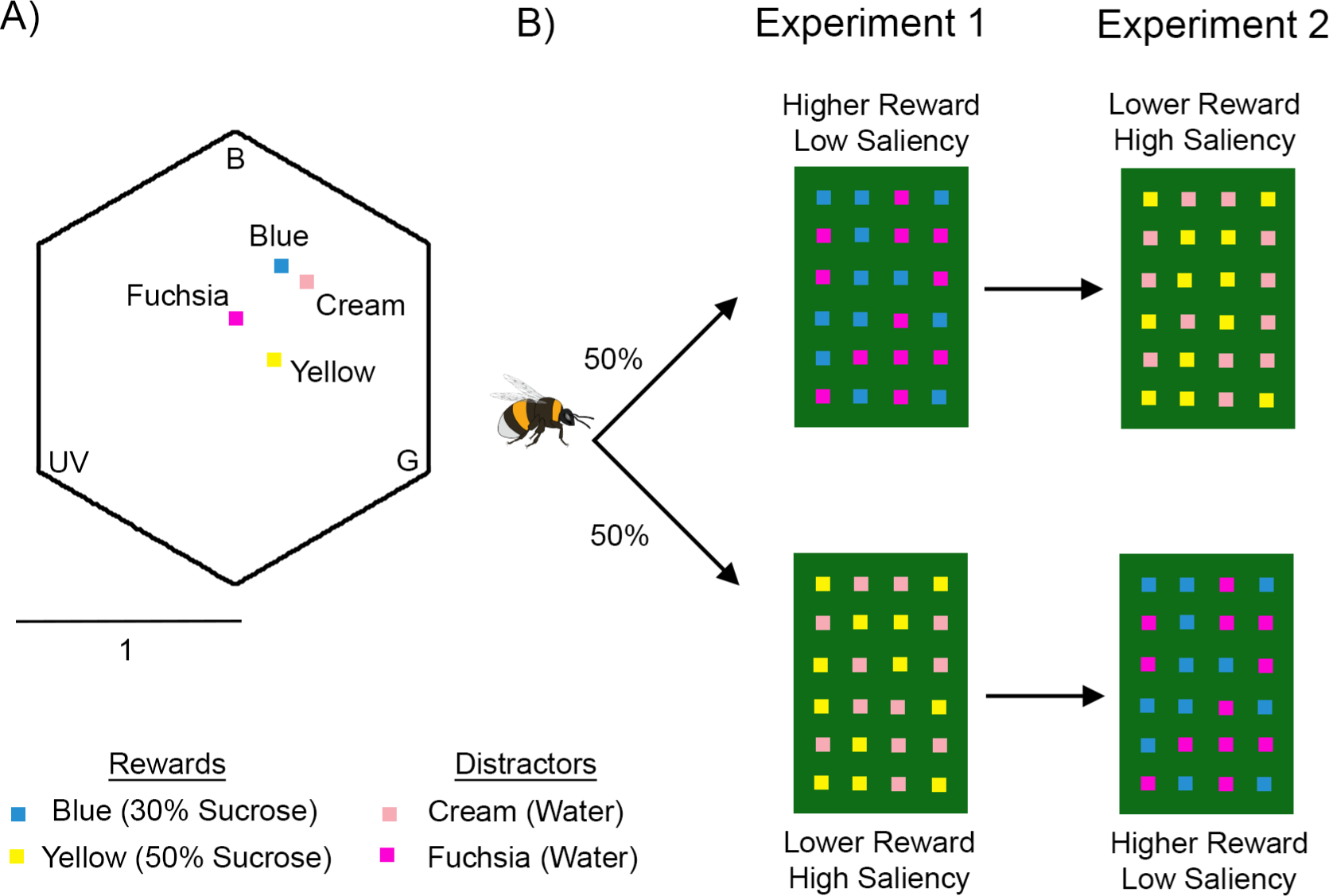
Experimental protocol. A) Colour loci of the artificial flower colours used in the experiments in the colour hexagon (Chittka 1992). Three vertices correspond to maximum excitation of photoreceptors sensitive to blue (B), green (G) and ultraviolet (UV) light. The distance from the centre to any vertex is 1 (see scale) and represents how salient is a colour. The distance between points represents hue discriminability, with 0.1 being easily distinguishable. B) Training paradigm in the experiments. Half the bees followed the protocol in the top row and half the bees followed the one in the bottom row. Rewarding (Blue and Yellow) and distractor (Fucshia and Cream) colours were the same in both but the order in which they were encountered (i.e. Experiment 1 or 2) was reversed. The rewarding flowers had 12 μl of 30% (lower reward) or 50% (higher reward) sucrose solution while distractors had an equal quantity of distilled water.

### Pretraining

We trained bees with no experience of colour to forage for sucrose solution from transparent square Perspex chips (side: 25 mm, thickness: 5 mm). These served as artificial flowers (henceforth “flowers”) and the aim of the pretraining was to allow the bees to learn to forage from them. Each flower had a central well that could be loaded with rewarding or unrewarding solutions. Once bees learned to forage from these chips, we placed them on glass vials (1.5 cm diameter, 4 cm tall) and trained bees to forage from them. We arranged 24 such vials in a 6 × 4 horizontal grid, placed 15 cm apart. Twelve of the flowers had 12 μl of 50% (v/v) sucrose solution in them and the others were empty. We randomized the positions of rewarding and non-rewarding flowers. In the pretraining and in all experiments these positions were randomized using the random number generator function RAND() in Microsoft Excel®. We moved to the training phase once the bee had foraged on this grid for three bouts.

### Training

We trained 16 bees from three colonies on a visual discrimination task. Bees had to discriminate between rewarding flowers (targets) of one colour and distractor flowers of another colour. These flowers were coloured Perspex chips placed on glass vials in a grid as described above. In each experiment, there were a total of 12 rewarding flowers and 12 distractors. Rewarding flowers contained 12 μl of 50% (v/v) sucrose solution while distractors contained 12 μl of distilled water. Within one foraging bout, flowers were not refilled but bees were allowed to revisit flowers multiple times. Bees were allowed to forage over multiple bouts until they made 80% correct choices of the rewarding flowers in their last 20 choices. Choices were recorded when the bees landed on a flower and probed them for reward, including when bees revisited flowers. Between training bouts, we cleaned the chips with 99% ethanol to remove scent markings, and then with water to remove traces of ethanol. The positions of the rewarding flowers and distractors was then randomized again before the next bout.

We trained each bee in two consecutive experiments (Figure 1B). The first looked at how training affected the visual search of naive bees and the role of reward quality. The second experiment looked at how prior training (in the first experiment) affected subsequent visual search when different rewards were encountered.

#### Experiment 1: The Effects of Colour-Naïve Training

Bees were divided into two groups (Figure 1B). The first group was trained on blue rewarding flowers with a lower reward of 30% (v/v) sucrose solution and fuchsia distractors. The second group were trained on yellow rewarding flowers with a higher reward of 50% (v/v) sucrose solution with cream distractors. We trained the bees until they reached the success criterion defined above.

#### Experiment 2: The Effects of Prior Training

In the second experiment (Figure 1B, right column), we continued to train the bees from Experiment 1 above on a novel task. For this task, we swapped the training regimes described in Experiment 1. Bees that were trained on lower rewarding blue flowers in Experiment 1 were now trained on higher rewarding yellow flowers with cream distractors. Bees in the other group that were trained in Experiment 1 on higher rewarding yellow flowers were now trained on lower rewarding blue flowers with fuchsia distractors. Higher rewards were always 50% (v/v) sucrose solution and lower rewards were always 30% (v/v) sucrose solution. Distractors always only held distilled water.

The choices made by the bees were noted to determine the success criterion and all bouts were recorded using a Sony DCR-SR58E Handycam at 25 frames per second.

### Video Coding

We analysed the videos using the open-source program Tracker (V5.15, ©2020 Douglas Brown, physlets.org/tracker). We perspective corrected the videos and tracked the position of the bees in each frame. Frames in which the bee was not clearly visible because of light reflections or because it flew to a corner of the arena were marked as missing data and excluded from subsequent analysis. We used the tracked position of the bees to obtain maps of bee search behaviour. To exclude time spent visiting a flower we excluded frames on which the bee was within 1.77 cm from the centre of a flower (for both distractors and rewarding flowers). This distance corresponds to the diagonal length from the centre to any corner of the artificial flowers we used. The visual search area was thus the area of our arena, after excluding the flower areas. Within the visual search area, we defined flower inspection regions on our maps as between 1.77 cm and 5 cm from the flowers’ centre. All other regions in the visual search area were defined as non-flower regions. We summed the total number of frames that a bee spent in each region and converted this to a measure of inspection time by dividing by the frame rate of the videos.

To investigate the effect of learning we compared the change in inspection time as our measure of attention. With this measure, we tested how attention for rewarding flowers, distractors, and other regions differed between the first six choices the bees made and the last six choices. We made the same comparison for both experiments.

### Statistical Analysis

All analyses were run on R (version 4.2.1). We analysed the results using the glm and glmmTMB function of the glmmTMB package (Brooks et al. 2017) to run general linear and generalized linear mixed models. We assessed the fit of all our models using the DHARMa package (Hartig 2022).

To analyse the inspection time results, we calculated the proportion of time spent in each region (reward, distractor or other) compared the total visual search time. To control for the differing areas of each region we divided these proportions by the total area corresponding to each region. We then log-transformed the weighted proportions and used this as the dependent variable in a general linear model. We used the models to test the three-way interaction effect of learning stage (first or last six), rewarding flower colour (blue or yellow) and region (reward, distractor or other), with each of these predictors included as factors. We ran the same analysis separately for both the first and second experiments. For the second experiment, we had to exclude two data points where the proportion of frames for the distractor was 0 and therefore could not be log-transformed.

We also ran an analysis on the duration (total number of frames) over which bees made their first and last choices. To do this we used the glmmTMB function from the glmmTMB package to run av generalised linear mixed model with duration as the dependent variable and a negative binomial family. The independent variables were learning stage and rewarding flower colour and bee identity was included as a random variable.

Finally, we analyzed the choices of the bees using a generalized linear mixed model with the proportion of choices of rewarding flowers as the dependent variable, bee identity as a random variable and a binomial family distribution and a logit link function. In consecutive models, we included rewarding flower colour, experiment and learning stage as independent variables and selected the best model after comparing the models with anova() function in R. Details of the model selection process are provided in the code in the supplementary material.

## Results

### Experiment 1

Our model shows a significant main effect of training stage on bee inspection time (GLM, Estimate = 0.957, S.E.= 0.350, P = 0.008; Figure 2A and B), showing that as bees learnt about the rewards they were more likely to spend time inspecting rewarding flowers. In the first training stage, there were no significant main effect of flowers of different reward value (GLM, Estimate = 0.132, S.E.= 0.350, P = 0.708, Figure 2A and B, pink plots). In this stage, bees were not significantly more likely to inspect rewarding flowers compared to distractors for both values of reward (Yellow flowers: GLM, Estimate = -0.022, S.E. = 0.350, P= 0.951; Blue flowers: GLM, Estimate = -0.264, S.E. = 0.350, P = 0.453, Figure 2B, pink plots). They were also not significantly more likely to inspect rewarding flowers compared to non-flower regions for either reward value (Yellow flowers: GLM, Estimate = 0.102, S.E. = 0.350, P = 0.771; Blue flowers: GLM, Estimate = -0.105, S.E. = 0.350, P= 0.765, Figure 2A, pink plots).

**Figure 2.**
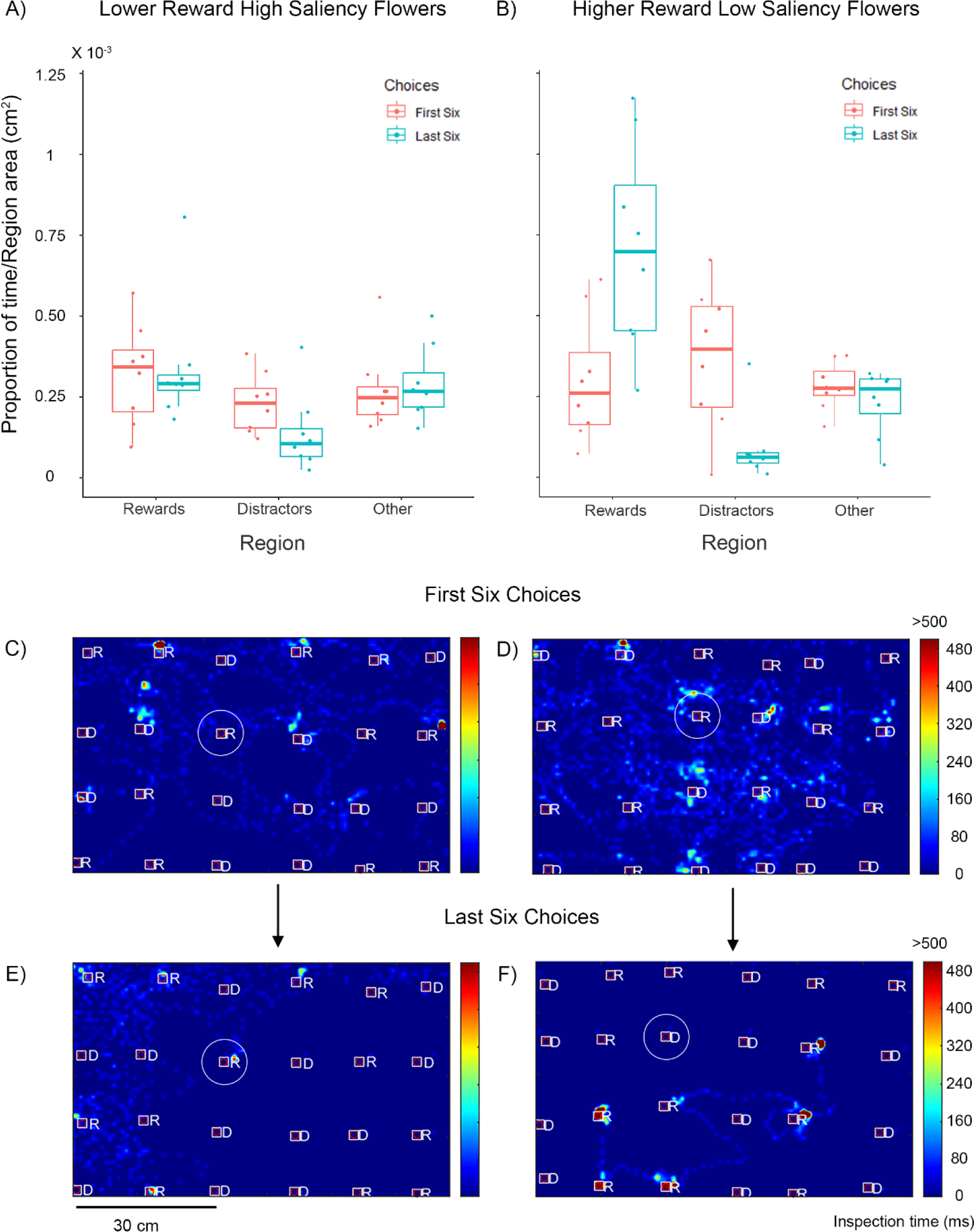
Bee visual search in Experiment 1. Top row: Proportion of time spent by the bees in different regions weighted by the area of each region. Bees were presented with A) lower reward (30% sucrose) high saliency blue flowers or B) higher reward (50% sucrose) lower saliency yellow flowers. Pink plots depict data from the first six choices and blue plots depict data from the last six choices of the training. Box plots depict the median and the first and third quartiles, the whiskers depict the largest and smallest values that are within 1.5 times the interquartile range from the edge of the boxes. Dots represent data from individual bees. C-F) Example visual search maps for two bees depicted as a top view of the flight arena with rewarding and distractor flowers. Colours depict the inspection times up to a maximum of 500 ms. Squares depict flower zones and the inner bound of the defined inspection zones, white circles illustrate the outer bound of the inspection zones. Only one circle is depicted in each figure here for ease of illustration. *R* = Rewarding flowers; *D* = Distractor flowers. C) and D) depict examples for the first six choices of training for bees trained on lower reward and higher reward flowers respectively. E) and F) depict the visual search during the last six choices of the same two bees as in C) and D) respectively.

For the higher rewarding yellow flowers, there were interaction effects between the training stage and the region type, showing that bees spent significantly more time inspecting these rewarding flowers compared to distractors and other regions in the later stage of training (Distractors: GLM, Estimate = -2.335, S.E.= 0.495, P < 0.001; Non-flower regions: GLM, Estimate = -1.282, S.E.= 0.495, P = 0.011; Figure 2A and B, blue plots). This was not true for the lower rewarding blue flowers (Distractors: GLM, Estimate = -0.843, S.E. = 0.495, P = 0.092; Non-flower regions: GLM, Estimate = - 0.019, S.E. = 0.495, P = 0.970; Figure 2A and B, blue plots)

In the later training stage, there was a main effect of reward value on the inspection time of reward, indicating that bees were more likely to inspect the higher rewarding yellow flowers compared to lower rewarding blue flowers (GLM, Estimate = -0.730, S.E. = 0.350, P = 0.040). At this training stage, bees inspected both types of flowers significantly more than distractors (Yellow flowers: GLM, Estimate = -2.356, S.E. = -6.738, P < 0.001; Blue flowers: Estimate = -1.107, S.E. = 0.350, P = 0.002). Bees trained on the yellow flowers, also inspected these higher rewarding flowers significantly more than non-flower regions (GLM, Estimate = -1.180, S.E. = 0.350, P = 0.001). Crucially, this was not true for bees trained on the lower rewarding blue flowers (GLM, Estimate = -0.124, S.E. = 0.350, P = 0.724).

These results suggest that bee attention to rewards is increased as they learn about the flowers. They also show that higher rewarding and lower rewarding flowers have slightly different effects. When flowers have lower rewards, bumblebees continue their searching behaviour rather than focussing on the rewarding flowers.

The number of correct choices made by the bees was best explained by a model that included the training stage and the flower (and thus reward) type as predictors but not the experiment (first or second). Bees were significantly less likely to make correct choices for blue flowers compared to yellow flowers (GLMM, Estimate = -1.525, S.E. = 0.464, P = 0.001). This likely reflects innate biases of the bees to blue but the bees were generally highly accurate. In the initial training stage of Experiment 1, bees made 79% correct choices to yellow flowers and 88% correct choices to blue flowers. The accuracy of the very first choices was however 50% (4 out of 8 bees) for yellow flowers and 62.5% (5 out of 8 bees) for blue flowers. These proportions were not different from chance (binomial tests, Yellow: P = 0.273, Blue: P = 0.219). In the later training stage, all bees were 100% accurate. This result also highlights the differences found when analysing choices and visual search and how both analyses complement each other.

Bees also made their last six choices faster than they made their first six choices (comparison with null model: χ^2^ = 26.216, df = 1, P < 0.001: First six VS last six: Estimate = -1.004, S.E. = 0.154, Z = -6.51, P < 0.001). Including flower colour as a factor did not improve the model (comparison with model having only training stage: χ^2^ = 0.753, df = 2, P = 0.686), suggesting that the effect was comparable in both higher and lower reward flowers. Learning thus increases the speed of choices regardless of flower reward level.

### Experiment 2

Unlike in Experiment 1, we did not find a main effect of training stage in this Experiment indicating that the training did not here increase bee inspection of rewarding flowers (Yellow flowers: GLM, Estimate = 0.384, S.E. = 0.244, P = 0.119, Blue flowers: GLM, Estimate = -0.218, S.E. = 0.244, P = 0.374, Figure 3A and B). In the first training stage, the group of bees that had switched from lower to higher rewarding flowers between experiments were more likely to inspect rewarding flowers compared to the other group (GLM, Estimate = 0.586, S.E. = 0.244, P = 0.019, Figure 3A and B, pink plots). This demonstrates the effect of the prior experience.

**Figure 3.**
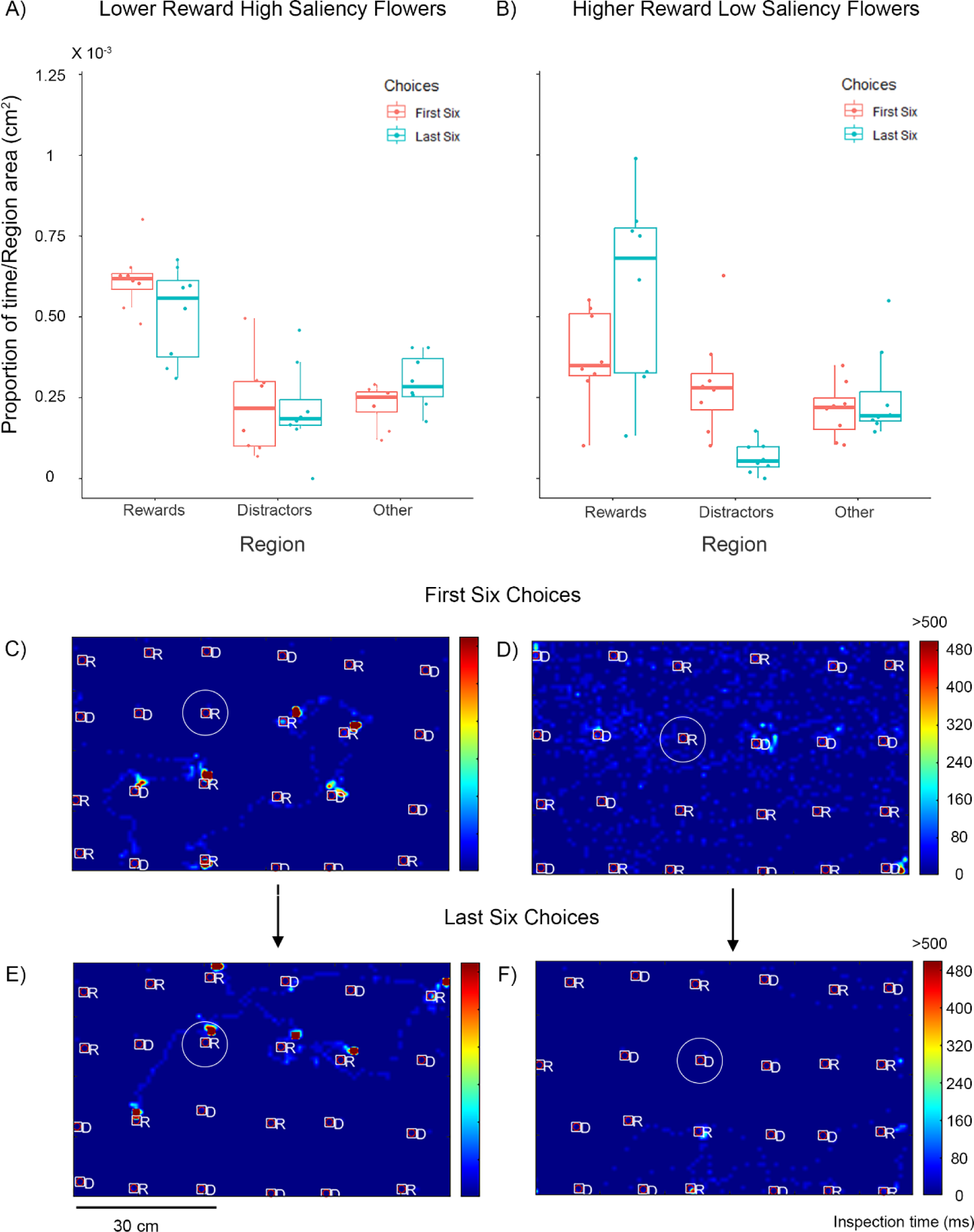
Bee visual search in Experiment 2. The data here represent bees initially trained on one reward type (higher or lower) in Experiment 1 (Figure 2) and subsequently trained on the opposite reward type (lower or higher) in Experiment 2. Top row: Proportion of time spent by the bees in different regions weighted by the area of each region. Bees were presented with A) lower reward (30% sucrose) high saliency blue flowers or B) higher reward (50% sucrose) lower saliency yellow flowers. Pink plots depict data from the first six choices and blue plots depict data from the last six choices of the training. Box plots depict the median and the first and third quartiles, the whiskers depict the largest and smallest values that are within 1.5 times the interquartile range from the edge of the boxes. Dots represent data from individual bees. C-F) Example visual search maps for two bees depicted as a top view of the flight arena with rewarding and distractor flowers. Colours depict the inspection times up to a maximum of 500 ms. Squares depict flower zones and the inner bound of the defined inspection zones, white circles illustrate the outer bound of the inspection zones. Only one circle is depicted in each figure here for ease of illustration. *R* = Rewarding flowers; *D* = Distractor flowers. C) and D) depict examples for the first six choices of training for bees trained on lower reward and higher reward flowers respectively. E) and F) depict the visual search during the last six choices of the same two bees as in C) and D) respectively. The example in C) and E) here is the same bee trained in D) and F) in Figure 2 and the example in D) and F) here is the same bee trained in C) and F) in Figure 2.

In this training stage, bees in both groups were significantly more likely to inspect rewarding flowers compared to non-flower regions (Yellow flowers: GLM, Estimate = -0.547, S.E. = 0.244, P = 0.028, Blue Flowers: GLM, Estimate = -1.02, S.E. = 0.244, P < 0.001, Figure 3A and B, pink plots). Bees with prior experience of high rewards also were significantly more likely to attend to the now lower rewarding flowers compared to distractors (GLM, Estimate = -1.197, S.E. = 0.244, P < 0.001). For bees in the other group this effect was not significant (GLM, Estimate = -0.272, S.E. = 0.244, P = 0.267).

There was a significant interaction effect showing that training increased attention to rewarding flowers compared to distractors when bees encountered higher rewarding yellow flowers (GLM, Estimate = -1.819, S.E. = 0.351, P < 0.001). This was not true for the lower rewarding blue flowers (GLM, Estimate = 0.423, S.E. = 0.351, P= 0.231). In the later training stage, there was no main effect of reward value (GLM, Estimate = -0.002. S. E. = 0.317, P = 0.996). In this stage, bees inspected rewarding flowers significantly more than distractors (Yellow flowers: GLM, Estimate = -2.063, S.E. = - 0.325, P < 0.001, Blue flowers: GLM, Estimate = -0.802, S.E. = 0.327, P = 0.017). They also inspected rewarding flowers significantly more than non-flower regions (Yellow flowers: GLM, Estimate = -0.854, S.E. = 0.250, P = 0.001; Blue flowers: Estimate = -0.583, S.E. = 0.255, P = 0.025).

Training thus boosted attention to rewarding flowers for the bees that had previously encountered low rewarding flowers in Experiment 1 but now encountered higher rewarding flowers. This confirms our predictions of the effect of prior experience on visual search to subsequent rewards. However, our prediction was not true for bees that switched from higher to lower rewards. Here bees continued attending to the rewarding flowers even though they now encountered lower rewards.

The choices of the bees in Experiment 2 were again highly accurate. In the initial training stage of Experiment 2, bees made 65% correct choices to yellow flowers and 92% correct choices to blue flowers. In the later stage, bees from both groups chose the rewarding flower with 100% accuracy. Contrary to what we observed in the first experiment, including flower colour made for a better model, suggesting that the effect of learning on the time taken to perform six choices differed with the reward quality (or colour) (Comparison with model including only training stage: χ^2^ = 7.544, df = 2, P = 0.023). There was no effect of training for bees that switched from higher rewarding flowers in Experiment 1 to lower rewarding flowers in Experiment 2 (First six blue vs last six blue: Estimate = - 0.212, S.E. = 0.156, Z = -1.36, P = 0.175). However, when the bees changed from lower rewarding flowers to higher rewarding flowers, their first six choices were faster than these of the other flower colour group (First six blue VS First six yellow: Estimate = 0.439, S.E. = 0.196, Z = 2.24, P = 0.025) and training further increased the speed at which these bees made their last six choices (First six yellow VS last six yellow: Estimate = -0.630, S.E. = 0.222, Z = -2.84, P = 0.005).

## Discussion

We tested whether learning modified visual search in two related experiments. We found that learning resulted in an increase in the proportion of time spent by the bees around rewarding flowers compared to distractors and notably to non-flower regions, but this depended on reward value and prior experience of rewards. Lower rewarding flowers led to greater visual search in areas away from both rewarding and distracting flowers. This suggests that attention is more widely distributed for lower rewarding flowers compared to higher rewarding flowers. It’s also important to note that since the distractors were not rewarding, they would also provide the bees with negative reinforcement. This would also partially explain the clear difference in attention between rewards and distractors. Bees with prior experience of higher reward were also more likely to persist in attending to rewarding flowers even when they later encountered lower rewards. In our experiments, we cannot disentangle the effect of reward quality and colour or saliency. However, the fact that the high saliency of low reward flowers did not increase inspection of these flowers in Experiment 1 indicates that our results are likely to reflect reward quality of the flowers rather than their saliency.

In primates, eye movements are often used as proxies for overt attention (Schütz et al. 2011). The duration spent looking at aspects of a scene have also been used to compute attentional maps (Henderson and Hayes 2018). Our maps of inspection time perhaps best parallel these attentional maps. Results from these studies of eye movements have shown that attention can be influenced by several factors including saliency, reward value and the structure of a scene (Navalpakkam et al. 2010; Schütz et al. 2011; Henderson and Hayes 2018). In non-primates, attentional limitations have most often been studied in predators (Dukas and Kamil 2000; Dukas et al. 2002; Dukas 2004). There, findings show that attentional resources are more focussed when searching for cryptic prey. Conversely, hunger leads to praying mantises widening their search for possible prey (Bertsch et al. 2019; Pickard et al. 2021). Our findings further argue that learning about reward value also influences attention in bees, even when the rewarding flowers are not cryptic.

The median nectar sugar concentrations for flowers in Europe and globally is around 40% (Pamminger et al. 2019). Our reward values therefore correspond to a higher than median reward (50%) and lower than median reward (30%). The latter corresponds to the reward value of flowers in the 25^th^ percentile. However, bees in our experiment were pretrained on 50% sucrose solution. This could have had some influence our results. Prior experience of a particular reward can influence future behaviour of bees and ants in response to higher or lower rewards – a phenomenon called incentive contrast (Bitterman 1975; Wendt et al. 2019). However, in our first experiment, we do not see a difference between visual search to the two rewards in the first training stage, suggesting that the experience of higher rewards during pre-training did not have an immediate effect. We do, however, see an increase in visual search to the rewards compared to other areas, but only in the later training stage, suggesting that as bees encounter higher (but not lower) rewards, they spend less time searching and more time inspecting rewarding flowers. We find some evidence that prior experience can influence visual search behaviour in line with ideas about incentive contrast – bees that experience lower rewards in Experiment 1 had increased attention to higher rewards during Experiment 2 and made faster choices. However, we did not find the converse for bees that switched from higher rewards to lower rewards. One confounding factor here could be that the lower rewarding blue flowers in Experiment 2 were more attractive due to the innate biases of bees. Alternatively, continuous experience of high rewards from the pre-training and Experiment 1 might have boosted the motivation of bees to a high level and persisted for a longer time.

Our results demonstrate the value of investigating search behaviour rather than focussing on flower choice alone. Previous work (Nityananda and Chittka 2021) has looked at bee visual search when faced with a choice between multiple rewarding flowers of different reward and saliency values. Research there found that reward value biased inspection time at flowers, even for lower saliency flowers. Other work has also shown that bees fly shorter distances after encountering rewarding flowers, compared to non-rewarding flowers (Dukas and Real 1993). Our results further show how reward value modifies bee visual search during learning. This difference in bee behaviour might specifically reflect foraging behaviour in bumblebees. Bumblebees are less flower constant than honeybees (Wells and Wells 1983; Waser 1986; Hill et al. 1997), sampling other flowers even when specializing on a specific flower type – behaviours that have been called ‘minoring’ and ‘majoring’ respectively (Heinrich 1976, 1979). Given that honeybees are more constant to flowers, it is possible that we might find different results from honeybees with more focussed attention even for lower rewarding flowers. Previous work on bee visual search has already shown differences between honeybees and bumblebees. Honeybees show serial visual search, while bumblebees are capable of parallel visual search – their visual search for a target is independent of the number of distractors (Spaethe et al. 2006; Morawetz and Spaethe 2012). Running similar experiments to ours with honeybees and other bees might bring up further interesting differences in visual search and attention.

Our findings suggest that flowers that have higher concentration of reward are more likely to have focussed attention from bumblebees. Bees that encounter flowers with lower rewards would be expected to keep searching even as they visit the flowers. We might therefore expect flowers with lower rewards to compensate for this loss of attention. One possibility is that these flowers might be more salient than flowers with higher rewards. However, blue flowers which are salient in temperate zones are actually the ones that are the most rewarding to bees (Giurfa et al. 1995). Another study in Australia found no significant correlation between reward values and chromatic contrast (Shrestha et al. 2020). Our results also show that simply having high saliency does not lead to focussed attention as the high saliency lower reward flowers did not attract greater bee visual search. Flowers with lower rewards might therefore be under selective pressure to either invest in multimodal cues (Kulahci et al. 2008) or include secondary compounds in their nectar that might affect pollinator memory and attention. Caffeine and nicotine have both been shown to have effects on bee learning (Wright et al. 2013; Couvillon et al. 2015; Baracchi et al. 2017; Arnold et al. 2021) and we would predict that they should be more likely to be present in the nectar of flowers with lower concentrations of reward.

Our findings demonstrate the importance of investigating bee behaviour beyond flower choices. Understanding how visual search and attention is influenced by a variety of factors could further enhance our understanding of pollination ecology and bee cognition.

## Data Availability

All code and data relevant to this paper are included as supplementary material to this paper.

## Acknowledgements

VN and TR are supported by a BBSRC David Phillips fellowship BB/S009760/1 to VN. This work was also partly supported by a Marie Curie Incoming International Fellowship (PIIF-GA-2009–253593) to VN.

## Author Contributions

VN conducted the experiment. KT and VN analysed the videos. VN and TR ran the statistical analyses and wrote the paper.

